# Bioinformatics Pipeline using JUDI: *Just Do It*

**DOI:** 10.1101/611764

**Authors:** Soumitra Pal, Teresa M. Przytycka

## Abstract

Large-scale data analysis in Bioinformatics requires executing several software in a pipelined fashion. Generally each stage in a pipeline takes considerable computing resources and several workflow management systems (WMS), e.g., Snakemake and Nextflow, have been developed to ensure in case of multiple invocation of the pipeline, only the bare minimum stages that are affected by the changes across invocations get executed. However, when the pipeline needs to be executed with different settings of parameters, e.g., thresholds, underlying algorithms, etc. these WMS require significant scripting to ensure an optimal execution of the pipeline. We developed *JUDI* on top of a Python based WMS, *DoIt*, for a systematic handling of pipeline parameter settings based on the principles of DBMS that simplifies plug-and-play scripting. The effectiveness of JUDI is demonstrated in a pipeline for analyzing large scale HT-SELEX data for transcription factor and DNA binding where JUDI reduces scripting by a factor of five.

**Availability:** https://github.com/ncbi/JUDI

## 1 Introduction

Large scale analysis of data in Bioinformatics requires many software to be executed in a pipelined fashion where output of a upstream stage is fed as input to a downstream stage. Generally each such software takes a considerable amount of computing resources such as CPU time and memory. If the pipeline needs to be executed multiple times, may be due to input or parameter variations, it is desired that only the bare minimum stages that are affected by the variations are re-executed. The rest should be reused from the previous execution. To address such requirement, many workflow management systems (WMS) have been developed. One of the earliest and widely used system is Make (Stallman *et al.*, 2004). Each stage of the pipeline is specified as a rule of commands in Make to generate the output(s) from the input(s). Make automatically figures out the interdependency among the rules and by using a directed acyclic graph (DAG) of the rules executes the commands in a suitable order such that all inputs are already available before a command is executed. Moreover, if all inputs are unchanged, Make skips the execution of that command.

Make has a steep learning curve, specifically for the biologists. Several newer WMS have been developed to specify Make rules in simpler languages, e.g., Snakemake (Köster and Rahmann, 2012) and Nextflow (Di Tommaso *et al.*, 2017) use simplified Python and Groovy, respectively. These WMS can also schedule the pipeline tasks efficiently on multiprocessor environments by exploiting the DAG of rules.

However, these WMS require significant effort in scripting to ensure optimal execution of the stages when the pipeline needs to be executed multiple times due to changes in different parameters such as thresholds, algorithmic methods, and hyperparameters of machine learning algorithms. Generally these changes affect only a part of the pipeline leaving some scope for resource saving by fresh execution of the affected parts only. Though Snakemake and Nextflow enable adhoc use of wildchar parameters for easy scripting, a systematic handling of parameters is lacking in the literature. Moreover, these systems focus on each stage of the pipeline separately and the lack of information sharing across stages makes plug-and-play of stages difficult. For example, if a file generated by one stage is used as input in multiple stages downstream then the details of the file need to be re-specified unnecessarily in each downstream stage making it tedious when there are several parameters.

We develop *JUDI* on top of a Python based build system, *DoIt* (Schettino, 2008), to systematically handle the issue of parameter settings using the principles of DBMS. By abstracting each file and task in a pipeline stage with an associated parameter database JUDI additionally enables true plug-and-play of pipeline stages. The novel ideas in JUDI not only simplify pipeline scripting but also reduce script size significantly.

## 2 JUDI

The basic unit of execution in DoIt (Schettino, 2008), the underlying workflow management in JUDI, is a task roughly equivalent to a collection of rules in the Make (Stallman et *al.*, 2004). Like other workflow systems JUDI uses the dependency among tasks based on their input and output files. However, the novelty of JUDI is a consolidated way of capturing the variability under which the pipeline being build could possibly be executed. This variability could be the parameters to the software used in the pipeline stages or could be some new parameters introduced for the overall pipeline. JUDI stores this variability in a simple table which contains one column for each parameter where each row indicates a value of the parameter. The database is populated one parameter at a time by taking a Cartesian product of the list of categorical values the parameter can take with the database constructed so far. Multiple parameters are populated through a table which need not be the Cartesian product of the columns. For example, consider parameter database PDA with two parameters: 1) *sample* ∈ {100, 101, 102, 103} representing sequencing data from four individuals, and 2) *group* ∈ {1, 2} representing the end of paired-end sequencing data, with overall 8 rows representing the Cartesian product of the two parameters.

Each JUDI file is associated with a parameter database and hence does in fact represent a collection of physical files each corresponding to one row of the parameter database. When associated with example database PDA, a JUDI file *reads* could represent the eight FASTQ *file instances*: `{100_1.fq, 100_2.fq, 101_1.fq, 101_2.fq, 102_1.fq, 102_2.fq, 103_1.fq, 103_2.fq}`. The mapping from the rows of the parameter database to the physical path of the file instances is stored by an extra column for path in the parameter database.

Each JUDI task has four components: 1) *inputs*, 2) *targets*, 3) *actions* and 4) *parameter database* where the first three components are similar to those in DoIt with the difference that each element of inputs and targets is a JUDI file instead of a physical file. Like a JUDI file, a JUDI task represents a collection of *task instances* each corresponding to a row of the parameter database. For example, a JUDI task with parameter database PDA, an input ‘reads’ of the example above and a target ‘bam’ could align each of the eight FASTQ files to a reference genome and generate eight BAM files where the alignment software and its arguments are specified through one of the actions for the task.

Consider a JUDI task *T* with parameter database *D_T_* and a JUDI file *F* (input or target) with parameter database *D_F_* with common parameters *X*. In many cases like the example above *D_F_* = *D_T_*. Otherwise, there could be four scenarios. First, when *D_T_* has an extra parameter *Y*, for each *y* ∈ *Y* instance (*x, y*) of *T* gets the same instance *x* of *F*, ∀*x* ∈ *X*. Second, when *D_F_* has an extra parameter *Y* instance *x* of *T* gets a list of all instances {∀*y* ∪ (*x, y*)} of *F*. Third, when *D_F_*, *D_T_* have same parameters but row *x* of *D_F_* is missing in *D_T_*, it does not create a problem as far as task instances are concerned. Final, when row *x* of *D_T_* is missing in *D_F_* similar to third case, the instance *x* of *T* is undefined and hence should be dealt as an error. Thus, to determine the instances of *F* accessed in an instance of *T* the following sequence of basic database operations are performed. Let *D_T,F_* denote the *left outer join* between *D_T_* and *D_F_* on common columns *X* and for each (unique) row *r* of *D_T_*, let 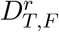 denote the projection Π*_r_* (*D_T,F_*). Then instance *r* of *T* gets all file instances in 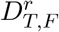. Along with the physical path information, 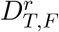 also contains the settings of extra parameters (if any) in *D_F_* and could be used by any summarization task as illustrated by an example below.

JUDI is available as a Python library and any JUDI pipeline first populates a global parameter database using the function `add_param` to avoid repeated local definition in each task. A task is a class derived from the base class `Task` and should have four class variables: 1) `mask`, 2) `inputs`, 3) `outputs`, 4) `actions`. Each of `inputs` and `outputs` is a python dictionary where each key names an input or target terminal of the task and has as value an object of JUDI class `File`. The object can be newly created using the class constructor or can refer to a File object created before. For example, if an input terminal of a task `A` is connected with to the target terminal `t` of another task `B`, then `A` can accessed the corresponding file(s) directly `B[t]`. In this way, the referencing task need not repeat the definition of the file.

If the class variable `mask` is set in a task then it represents the set of parameters from the global parameters database that are not applicable to the current task and the masked parameter database for the current task is created accordingly. Otherwise the task parameter database is assumed to be the same as the global parameter database. The actions for a task are specified by the python list `actions` of tuples (*fun, args*) where *fun* could be a Python string denoting the command line specification of the action with placeholders {} which are replaced by the list *args* of values similar to Python `format` function with following exception: if a value is “`$x`” then the placeholder is replaced by a string containing the list of paths of the file instances in 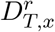 for the task instance *r*. A *fun* could also be a Python function and *args* could additionally have a value “`#x`” which JUDI replaces by a pandas DataFrame for 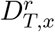 to pass to *fun*.

To illustrate the ideas used in JUDI, let us consider a slightly modified version of the 4-stage pipeline used in the Snakemake paper (Köster and Rahmann, 2012, see Fig. 1). In the first stage, each of the eight FASTQ files, one for each combination of 4 samples and 2 groups of pair-end reads is aligned to an intermediate file. The intermediate files for the two groups of pair-end reads for each sample is converted to a BAM file in the second stage. For each sample, the third stage generates a table containing genome coverage information. The fourth stage consolidates genome coverage of all samples into a single table and then plots this table, unlike the Snakemake example where the coverage plot is generated separately for each sample. The pipeline is shown in Fig. 1 where parameter databases are also shown along with each task and file. Listing 1 shows the Python script for the pipeline shown in Fig. 1. When command line `doit-f dodo.py` is executed, JUDI python library creates the task instances which are executed by the DoIt engine. More details of JUDI and DoIt can be found in their respective users manuals.

**Figure 1:**
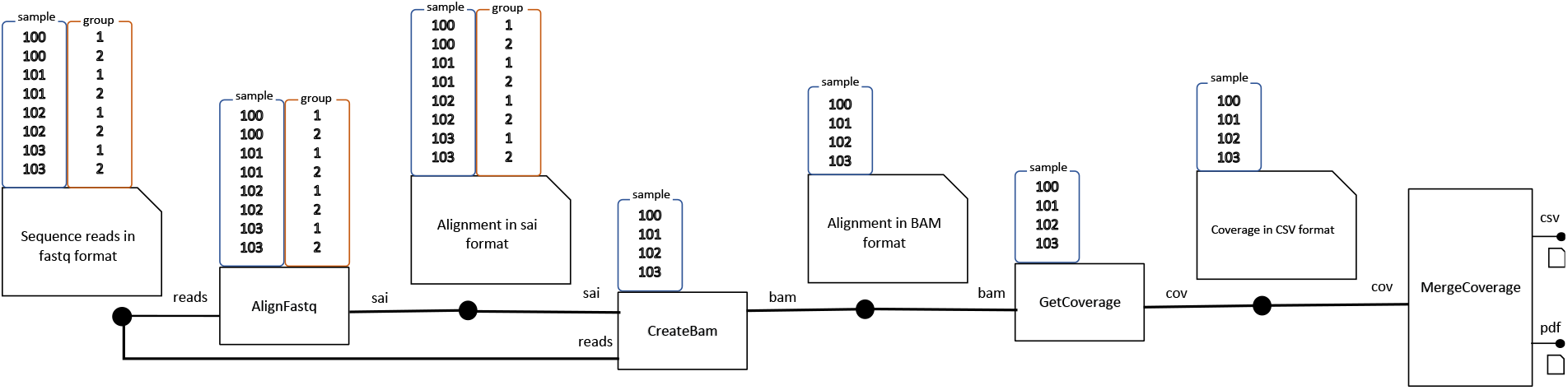
A slightly modified pipeline of Snakemake paper (Köster and Rahmann, 2012) visualized using JUDI concepts.

The usefulness of JUDI can be illustrated with two tasks in Listing 1. CreateBam has only one parameter ‘sample’ whereas its both input files ‘reads’ and ‘sai’ have both parameters ‘sample’ and ‘group’. For each sample, the physical paths of sai and fastq files for both pair-end reads are passed to the aligner `bwa sampe` automatically by JUDI through the argument substitutions `$sai` and `$reads`. Similarly, CombineCoverage has no parameter but its input ‘cov’ has a parameter ‘sample’. The JUDI library function `combine_csvs` merges the coverage files for all samples using 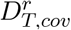 and 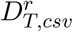 provided by argument substitutions `#cov` and `#csv`, respectively, where T denotes CombineCoverage.

## 3 A case study

We applied JUDI to the Co-SELECT tool (Pal *et al.*, 2018) developed for the analysis of HT-SELEX data for transcription factor (TF) DNA binding. The tool analyzed 5 *rounds* of data from 81 *TF experiments* for 3 *families* by dividing the sequencing reads into two *populations* to find statistically significant shape-strings which were enriched at 5 possible *thresholds* in both populations. Co-SELECT also analyzed *control experiments* taking cross population data for two TFs in two different families.

The implementation of the Co-SELECT pipeline took about 150 DoIt tasks with overall 2100 lines of Python code. On the other hand, these 158 tasks take less than 150×3 = 450 lines in JUDI. This gives about 5 times reduction in scripting. This significantly reduces the effort required to maintain the code. Moreover, the functions like `combine_csvs` further reduce the effort in summarizing results.

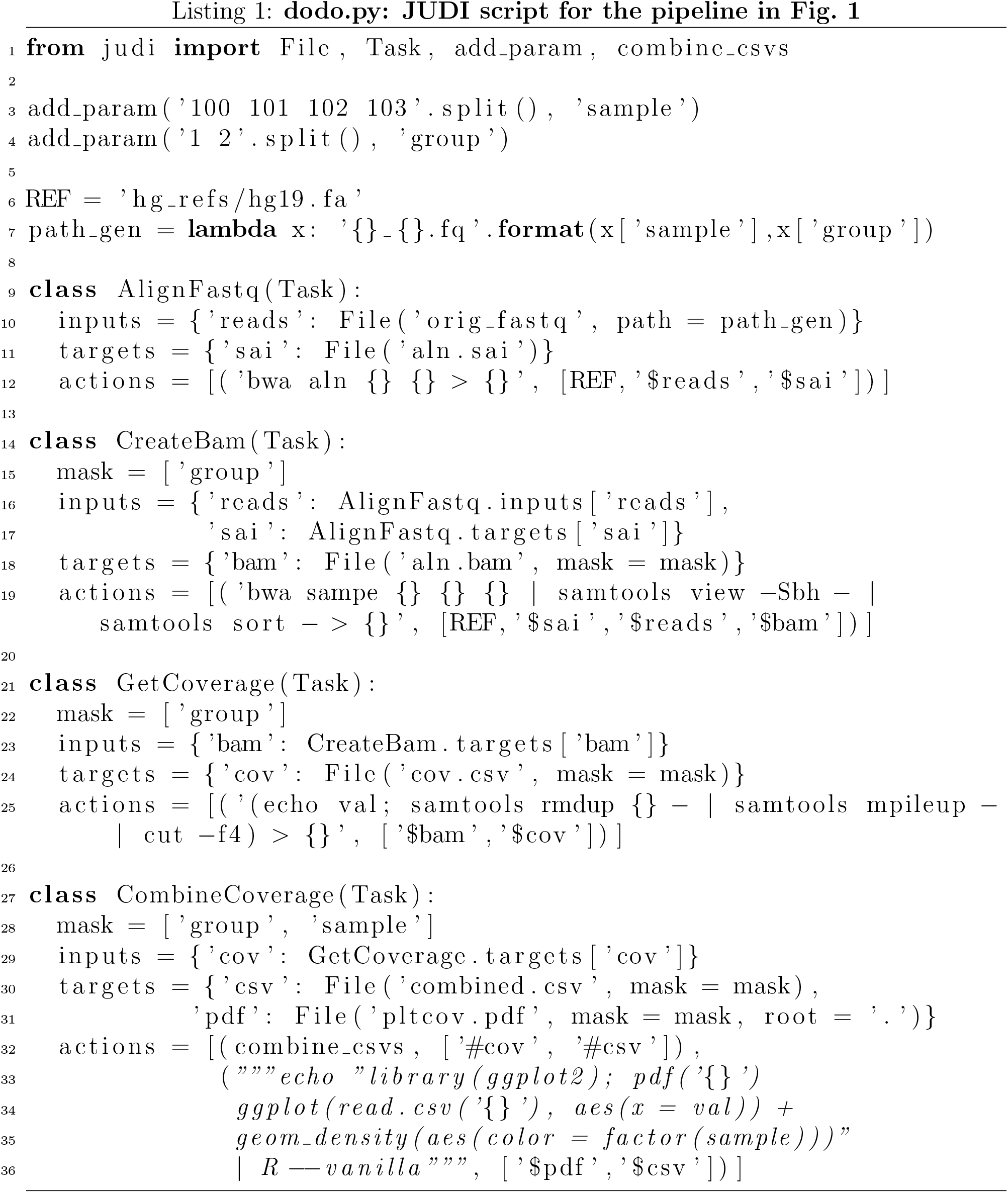

## 4 Discussion

In this paper, we introduced a workflow management system with a novel way of handling parameter settings and file path specifications with the motto: define once, reuse many times. We have implemented our ideas using an existing Python based build system DoIt (Schettino, 2008) mainly for two reasons: 1) DoIt does not require to learn any new language in addition to Python, and 2) it is more flexible for implementing our ideas quickly. However, our ideas can also be implemented in other WMS such as Snakemake (Köster and Rahmann, 2012) and Nextflow (Di Tommaso *et al.*, 2017). Though for the simple example in Fig. 1 JUDI takes almost same number of lines as Snakemake, for a larger pipeline as in Pal *et al.* (2018) JUDI requires far less scripting. For example, the file name expansion due to the masked parameter ‘group’ needed hard-coding in lines 11-12, Listing 1 of Köster and Rahmann (2012), imagine the effort required if there were more masked parameters and one or more parameters had a relatively large number of possible values!

There are a few features in Snakemake (Köster and Rahmann, 2012) and Nextflow (Di Tommaso *et al.*, 2017) which are not implemented in JUDI which could be easily implemented in a future version. One such example is to support temporary intermediate files which are not saved across invocation of the pipeline. Nextflow also provides an advanced way of handling files using stream, here we confined on file.

In the current implementation, the mask on the parameter database omits only the columns. However it will not be hard to implement one in which the mask omits some rows too.

Finally our solution can be extended to have graphical interface for better usability. When combined with interfaces like iPython Notebooks, JUDI could accelerate reproducible research.

## Acknowledgements

If JUDI has been convenient and more useful than other workflow systems, it is by standing upon the shoulders of giants like doit, we acknowledge their developers. This research was supported by the Intramural Research Program of the National Library of Medicine.

